# Detection of ‘*relict*’ western lineage of citrus tristeza virus virulent genotype in declining *Arunachal Wakro orange*

**DOI:** 10.1101/2020.08.18.255943

**Authors:** Sibnarayan Datta, Bidisha Das, Raghvendra Budhauliya, Reji Gopalakrishnan, Vanlalh Muaka, Mukesh K Meghvansi, Safior Rahman, Sanjai K Dwivedi, Vijay Veer

## Abstract

Eastern Himalayan foothills are known to have optimal agro-climatic conditions for production of quality citrus fruits including oranges. Among the citrus growing regions of eastern Himalayas, Wakro in the far eastern state of Arunachal Pradesh is known for its superior quality oranges, popular as the *Wakro orange* or the *Arunachal orange*, which has been included in Geographical Indication Registry by the Government of India. However, during last few years, Arunachal orange orchards have been experiencing severe infestation of aphid associated with rapid decline disease, causing catastrophe to the farmers as well as state economy. Therefore, in 2015, an intensive survey of severely affected orange orchards was carried out to investigate the aetiological factors of citrus decline. RNA samples extracted from leaf and aphid specimens collected from Wakro orchards were subjected to CTV detection through 3’-UTR specific RT-PCR. Subsequently, ORF1 and CP genetic regions were amplified and clonal-sequencing was performed. Although, BLAST search showed close homology of the present sequences with other virulent genotype VT sequences, detailed phylogenetic analysis demonstrated affinity and clustering of present sequences with VT sequences belonging to the ‘western’ lineage. This finding is considerably distinct from CTV sequences reported from citrus growing orchards in India and other neighbouring countries. Additionally, low diversity of CP gene sequences, recombination patterns and presence of sequence segments identical to the present ones in other CTV genotypes was also revealed. Collectively, these observations indicate pristine and primeval nature of present CTV sequences, corroborating well with the proposed origin of CTV in this part of the globe. We here report our finding of western lineage of CTV virulent genotype VT, which is distinct from CTV molecular epidemiology in other parts of India and discuss the implications of these findings.

## INTRODUCTION

Citrus group of fruits include oranges, mandarins, grapefruits, lemons, limes etc. and are widely grown throughout the world in large quantities^1^. However, production of citrus is largely restricted by the citrus tristeza virus (CTV), which is one of the economically most devastating viral pathogens of citrus. CTV is prevalent in almost all major citrus growing regions of the world and can infect almost all citrus species and hybrids and has caused a loss of more than 100 million citrus trees during the last century^2-5^. Long distance spread of CTV occurs via movement of vegetatively propagating material such as infected grafts (primary propagation), while local spread (secondary propagation) is facilitated by diverse aphid species in a semipersistent manner, of which the brown citrus aphid (BrCA, *Toxoptera citricida*) and the melon aphid *(Aphis gossypii)* are reported to be the most efficient vectors^2,3,6^. CTV is not known to be spread by seeds^3,4^.

CTV belongs to the genus *Closterovirus*, under family *Closteroviridae*, characterized by long flexuous rod-like virus particles (∼2,000x 12 nm) composed of two types of coat protein (major & minor coat proteins), encapsulating a single stranded, monopartite, linear, positive sense genomic RNA of approximately 19.3 kilobases^2,7,8^. The CTV gRNA consists of at least 12 open reading frames flanked by untranslated regions (UTRs) at both the 5’ and 3’ ends^2^. The ORF1a segment of the ORF1 encodes a large polyprotein having papain-like protease (P-Pro), Methyltransferase (Mt), and Helicase (Hel) activities, while the ORF1b encodes RNA dependent RNA polymerase (RdRp), expressed through +1 frameshift mechanism^2,6,8^. ORFs 2–11 encode at least ten distinct proteins, namely p33, p6, HSP70h, p61, CPm, CP, p18, p13, p20 and p23, expressed through subgenomic RNAs^2,6,8^. These structural and non-structural proteins are involved in capsid formation, virion assembly, suppression of plant RNA silencing mechanism, host range determination etc^2,6^.

As per available ICTV taxonomy report, three distinct strains of CTV have been identified, namely the T30 (mild isolate), T36 (intermediate severity) and VT (severe isolate)^8^. However, availability of complete genome sequences and detailed phylogenetic analyses has led to identification of at least eight different CTV genotypes distributed throughout the globe, namely the T36, T30, T3, VT, B165, HA16-5, T68 and the resistance breaking RB^9^. Genotype of infecting CTV is considered as an important factor that determines magnitude of damage^2,3^.

Citrus species are among the economically most important horticulture crops and are cultivated throughout the tropical and sub-tropical geographical zones of India^7,10^. Among them, mandarins (*Citrus reticulata* Blanco) particularly the *Nagpur mandarin* (Central India), *Kinnow mandarin* (Northwestern India), *Coorg mandarin* (Southern India*), Darjeeling mandarin, Sikkim mandarin* or *Khasi mandarin* (Northeastern India, NE India) and the *Arunachal Orange/ Wakro Orange*/ *Arunachal Wakro Orange* (NE India) are economically important cultivars. Among others, sub-tropical agro-climatic conditions of eastern Himalayan regions (including Darjeeling hills, Sikkim, Arunachal Pradesh) of NE India is known to be optimal for growing quality citrus fruits^11,12^. Similar to other citrus-growing regions, Indian citriculture is also severely threatened by CTV. CTV has killed estimated one million citrus trees in India during the last century^13^. Percentage of disease incidence varies widely throughout India^7,13-15^. Some NE Indian orchards in Assam and Meghalaya are reported to have even 100% incidence of CTV infection^16^. A number of previous studies have reported prevalence of genotype VT or its recombinants, with extensive diversity of coat protein (CP) gene in different citrus growing regions of India^17-20^.

Among the NE Indian State of Arunachal Pradesh (AP), citrus fruits represent the largest horticulture crop, of which the *Arunachal mandarin* or the *Arunachal Wakro orange* or *Wakro orange* (named after place of its origin – *Wakro*, 28.113°N and 96.812°E, Lohit district) accounts for 90% of total citrus production. *Wakro orange* is largely organic and has certain unique features that make it superior from other mandarin oranges grown in other parts of India. High vitamin-C and total soluble solid (TSS) content, medium acidity sweet-sour taste, loose and easily removable medium thick peel, good size, better juice content and an attractive orange colour of this variety is attributed to characteristic agro-climatic conditions of AP^21^. Owing to its distinctive characteristics, *Wakro Orange* has been registered under the *Geographical Indications of Goods (Registration and Protection) Act, 1999* by the Government of India^21^.

Unfortunately, during the last few years, Wakro orange has been witnessing a sharp decline in production, severely affecting the economy of orange growers as well as of the state^22^. A recent survey conducted by the Arunachal Pradesh Horticulture Research and Development Mission (APHRDM) found this “*Citrus Decline*” situation highly alarming, where farmers were even forced to abandon hundreds of hectares of orchards, completely engulfed by the rapidly spreading decline disease^23,24^. After survey, APHRDM officials suspected bacterial and/or viral infection, nutrient deficiency etc^23^. Despite presence of diverse aphid species including *T. aurantii and T. citricida*^21^ and indication of CTV infection in several quick declining citrus orchards^25^, molecular epidemiology of CTV in this part of India has not been studied systematically. This dearth of information may be largely attributable to location of *Wakro Orange* orchards in tough terrains in some of the remotest parts of India.

Our present study was carried out with an aim to detect, identify and characterize CTV associated with rapid decline of Wakro oranges in this region.

## MATERIALS AND METHODS

### Sample collection

During March 2015, a survey of severely declining orange orchards in the Wakro circle (also known as the ‘*orange bowl of the State*’) of the Lohit district in Arunachal Pradesh, India was conducted, based on request of district horticultural department. Sampling was done from affected orchards at four sites namely *Karhe* (27°54.20’N 96°17.38’E; 365 m ASL), *Cibi* (27°44.77’N 96°20.24’E; 345 m ASL), *Kangjang* (27°48.15’N 96°20.78’E; 460 m ASL), *Mawai* (27°48.15’N 96°20.75’E; 330 m ASL) and from one healthy Government maintained healthy orchard at *Mawai* (27°49.66’N 96°19.91’E; 326 m ASL). Location of the study site, and some glimpses of the affected orchards are shown in Figure 1. Leaf samples along with aphids were collected from affected plants, immediately preserved in RNAlater (Sigma-Aldrich, St. Louis, USA) and transported to laboratory for further investigations.

**Figure 1.**
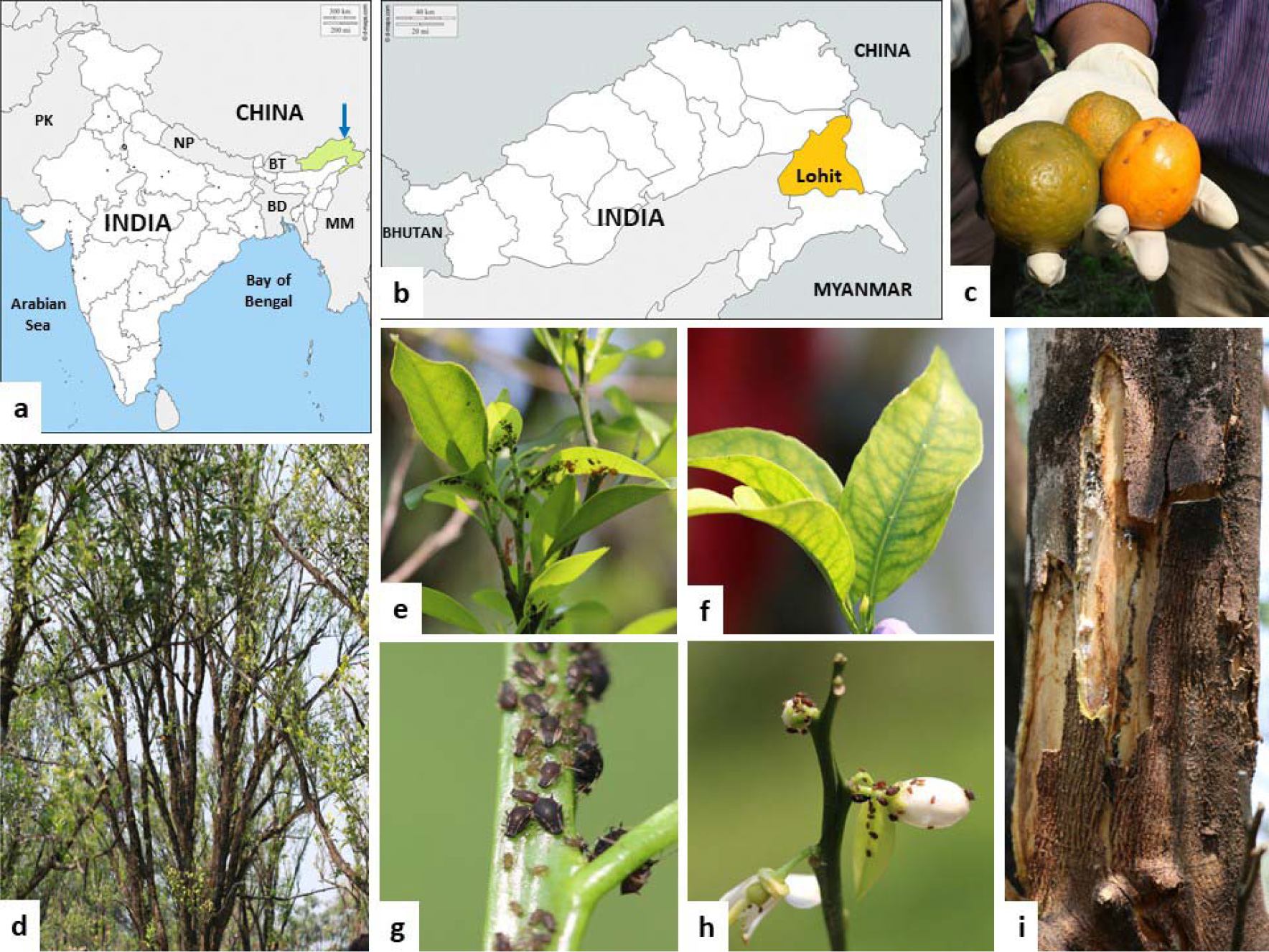
(a) Map showing the extreme northeast Indian state of Arunachal Pradesh (in light green and indicated by an arrowhead). Neighbouring countries are denoted by alpha-2 country codes: MM, Myanmar; BD, Bangladesh; BT, Bhutan; NP, Nepal; PK, Pakistan. (b) Map of Arunachal Pradesh showing the Lohit district (in orange colour). Panels (c) to (i) show fruits from diseased trees, affected orchards, aphid colonies on leaves and affected leaves with disease symptoms, observed during field survey. Blank base maps were obtained from the *d-maps* website (*https://d-maps.com/index.php?lang=en*) and annotated using Microsoft Office PowerPoint 2007 and Microsoft Windows Paint softwares.

### RNA extraction and detection of CTV RNA

Approximately 1 mg of individual leaf sample and aphid pools (each composed of 10-15 aphids) were ground in ball mill (Retsch GmbH, Düsseldorf, Germany) and total RNA was extracted from homogenate using standard TRI-reagent protocol (Sigma-Aldrich). Following assessment of quality and quantity (denaturing agarose gels and spectrophotometry), presence of enzyme inhibitors in RNA extracts was assessed by reverse transcription PCR (eAMV RT, Sigma-Aldrich) using universal primers for 18S RNA. Appropriate control reactions were included during each reaction.

Initially, CTV was detected using RT-PCR targeting amplification of highly conserved 3’-UTR of the CTV genome^26^. Briefly, first round of RT-PCR was done with external primer pairs PEX1/PEX2, followed by second round of amplification with internal primer pairs PIN1/PIN2, generating an amplicon of approximately 132 bps, which indicated presence of CTV genome. Specificity of the 132 bp amplicons was verified by restriction digestion with *Alu* I, which yielded two fragments of approximately 86 bp and 46 bp, as expected. Restriction digestion pattern of the amplicon was predicted using an alignment of 3’-UTR region from 70 CTV genomes belonging to various genetic types, available in the NCBI GenBank.

### Amplification, cloning and sequencing of CTV major CP and RdRp genetic regions

For amplification of CP gene sequence, we used previously described PCR primers-HCP_1_ and HCP_2_, which amplify complete CP ORF of 672 bps^27^. The ORF1 encoding complete RdRp ORF (RNA dependent RNA polymerase) was amplified using two pairs of overlapping primers, namely SY17/SY18 and SY19/SY20, described earlier^28^.

Following amplification, PCR products were purified, cloned into vectors using T/A cloning kit (Fermentas, Villinius, Lithuania) and were used to transform XL1-Blue competent bacterial cells. For each amplicon 5-10 clones were randomly selected, cultured and plasmid DNA purified (Sigma-Aldrich). Subsequently, plasmid DNA was amplified using vector specific universal primers (M13F and M13R) and sequenced (AgriGenome Labs Pvt Ltd., Kochi, India).

### Molecular evolutionary analysis

After trimming the vector sequences from sequence data, alignment, manual editing and assembly (for overlapping amplicons) were done using Bioedit version 7.2.5^29^. Sequence alignments were subsequently analysed through *Elimdupes* program (*https://www.hiv.lanl.gov/content/sequence/elimdupesv2/elimdupes.html*) to identify identical or near identical sequences among clonal sequences, based on inter-sequence percent nucleotide identity (PNI). From each group of identical or near identical sequences (PNI ≥ 98), one representative sequence was selected for further analyses and submission to the GenBank. Integrity of protein coding nucleotide sequences was verified through *ORFfinder* program (www.ncbi.nlm.nih.gov/orffinder/). Sequences were then BLAST (BLASTn, Megablast) searched^30^ to recognize and retrieve homologous sequences available in the GenBank non-redundant (nr) database. Additionally, well-characterized CTV sequences enlisted in the ICTV species list^8^ and reviewed in recent literature^2,7^ were also retrieved from GenBank for further analyses.

Sequences were aligned using MUSCLE algorithm^31^ followed by molecular evolutionary analyses using MEGA version 6^32^. Best fitting nucleotide substitution model was determined from lowest Bayesian Information Criterion (BIC) score. Evolutionary distances were calculated using Maximum Composite Likelihood method and phylogenetic relatedness was determined employing Neighbour-Joining (NJ) algorithm. To keep the phylogenetic tree uncluttered, among similar or nearly similar GenBank sequences, identical or near identical sequences arising from same site/time/study were kept to minimum. Consistencies of tree branches were computed by 1000 bootstrap replications of the datasets.

To avert ambiguities in sequence relatedness, which arise from recombination events, we reconstructed Neighbour Network (NeighborNet or NN), using *SplitsTree* version 4.14.1^33^, with Kimura-2 parameter distance correction algorithm. Data gaps and parsimony uninformative sites were excluded from the calculations. NN reconstruction algorithms accommodate recombination events and represent sequence relatedness in more accurate topologies^34^.

To examine evidence of recombination events in present sequences, aligned datasets were analysed through different recombination detection algorithms, namely RDP, GENECONV, BOOTSCAN, MAXCHI, CHIMAERA, SISCAN, and 3SEQ, incorporated in RDP version 4.72^35^. Computations were done on default settings with standard *Bonferroni* correction. Recombination events with *P<0*.*01* were further verified by analysis of breakpoint plots and phylogenetic trees (Unweighted Pair Group Method with arithmetic Mean; UPGMA) generated by the program.

## RESULTS

### Detection, amplification, cloning and sequencing of CTV gRNA from Citrus plants

Initially, leaf RNA extracts from five different orchards (4 affected and 1 healthy) and RNA extracts from four aphid pools (collected from affected orchards) were subjected to 3’-UTR RT-PCR. Amplicons of expected size were repeatedly detectable in leaf RNA extracts from affected orchards, but not in extracts from healthy orchard (Figure 2A). Additionally, amplicons were also detected in three of four aphid pools, collected from different declining orchards. Specificity of the 132 bp amplicons was confirmed by digestion with restriction endonuclease *AluI*, which yielded two fragments of 86 bp and 46 bp (Figure 2B), and also by direct sequencing of the 132 bp amplicons. Complete CP and RdRp coding genetic regions could be reproducibly amplified in three of four leaf RNA extracts that were positive by 3’-UTR RT-PCR. Nevertheless, despite repeated attempts, we could not amplify these genetic regions in one of the 3’UTR RT-PCR positive leaf RNA extracts or in any of the aphid pool extracts.

**Figure 2.**
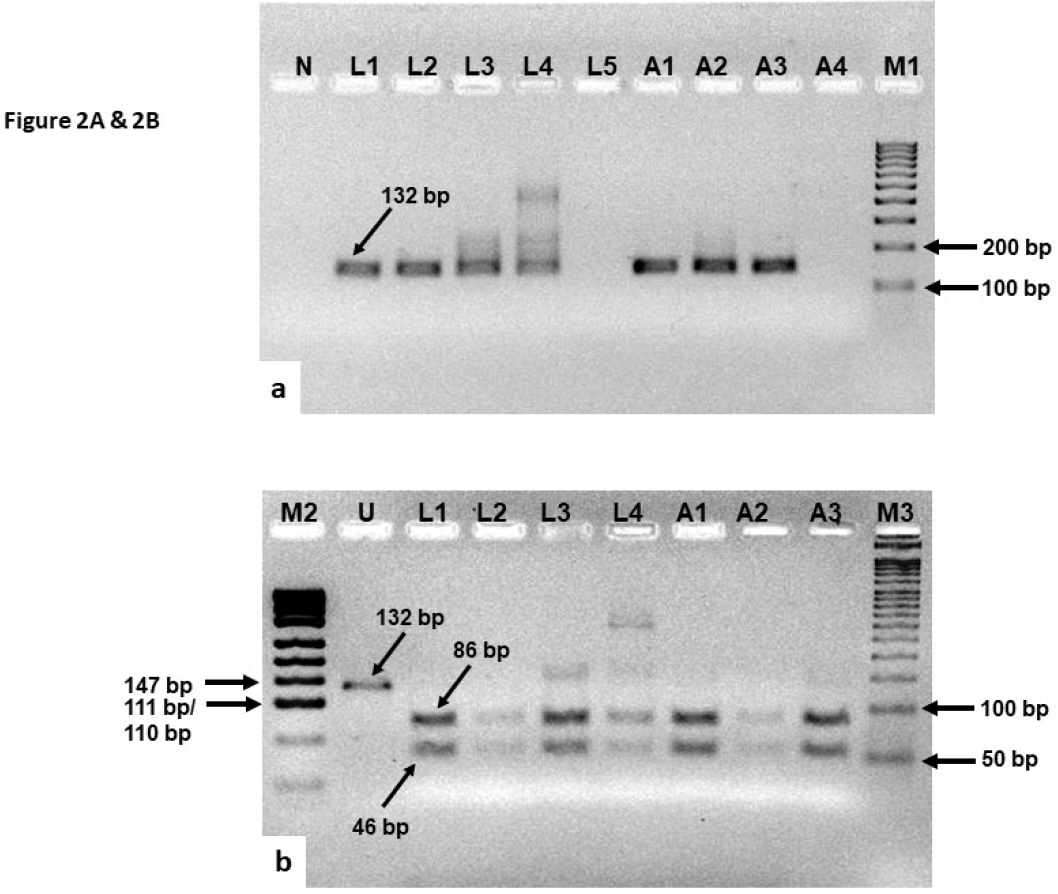
(A) Agarose gel showing 132 bp amplicon from CTV 3’-UTR RT-PCR. Lanes N, No Template Control (NTC); L1 to L5, RNA from leaf extracts; A1 to A3, RNA extracts from aphid pools. Marker, M1 is 100 bp DNA ladder. (B) Agarose gel showing *Alu* I digested products of 132 bp amplicons. Lanes U, uncut amplicon; L1 to L4, amplicons from leaf extracts; A1 to A3, amplicons from aphid pools. Marker, M2 is pUC19 DNA/MspI digested; M3 is 50 bp DNA ladder. Gel images were acquired on an UVP BioSpectrum 410 imaging system (UVP, Cambridge, UK) with automatic exposure setting. Colour of the original gel image was inverted using Microsoft Windows Paint and annotated using Microsoft Office PowerPoint 2007.

Subsequent to clonal-sequencing of the CP and RdRp amplicons, alignment of homologous sequences and screening through *Elimdupes* program, one representative sequence from each group of identical clonal-sequences was selected for further analyses. Finally, three representative CP gene sequences (S21, S24 & S28) and two representative RdRp gene sequences (S5 & S9) were analyzed in details.

Nucleotide sequences generated in this study are available in the NCBI GenBank under accession numbers - KY882459 through KY882476.

### Diversity of the sequences and results of BLAST analysis

Among the complete CP gene sequences, S21 and S24 were found to be virtually identical, sharing a PNI of 99% among themselves. While sequence S28 was significantly divergent from S21 and S24, sharing a PNI score of only 93% (S28 vs. S21 & S24, P<0.01). On the other hand, the two representative RdRp sequences S5 and S9 shared a PNI of 98% among themselves.

In BLASTn search, CP sequences S21 and S24 were found to be highly homologous (99% identity at 100% query coverage) to a number of sequences isolated from different parts of India as well as other countries, viz. eastern and NE India (HM573451, GQ392063, GQ475551-GQ475553, LT576375), western India (GQ475563, GQ475564), Southern India (EU869297), Northern India (JN974902, FJ001829), Uruguay (KU900351), Argentina (EU579381), South Africa (KU883267), Reunion Islands (AY660010) and Malaysia (HQ012413). Conversely, sequence S28 was found to be highly homologous (99% identity at 100% query coverage) to sequences recently isolated from different parts of NE India (GQ475549, KR185332; KR185331, KR185329, KR105771; KR259640, KR259641, KR080487, LN997804) as well as from New Zealand (AY896563).

On the other hand, in BLASTn analysis, RdRp sequences-S5 and S9 were found to be highly similar (97-98% identity at 100% query coverage) to CTV genotype VT sequences from USA (KC517493, EU937519, KC517494; KC517492; KU361339), Greece (KC262793), China (JQ911664, KU720382, JQ061137), Spain (DQ151548), Italy (KJ790175, KC748392), Japan (AB046398) and Israel (U56902). Interestingly, present sequences were reasonably divergent from two distinct complete genome CTV sequences isolated from India, namely the Kpg3 from Eastern India (HM573451, 94% identity at 100% query coverage) and the B-165 from southern India (EU076703; 86% identity at 100% query coverage).

### Phylogenetic affinities of the CP and RdRp genetic regions

In the CP gene phylogenetic tree, sequences from various parts of India and other countries segregated into eight distinct clades previously classified as genogroups I through VIII^36^. Clustering of present sequences S21 and S24 corroborated with BLASTn results, demonstrating their affinity with group VII of CP gene sequences (Figure 3). Conversely, sequence S28 along with two other sequences (AY896563 from New Zealand & KR259640 from Sikkim, NE India) acquired an intermediate position between group I sequence clade (which comprise sequences mostly isolated from NE India) and group VIII sequence (KC590504 from Tirupati, south India) (Figure 3). This intermediate position of sequence S28 was more prominent in NN analysis (Figure 4). Although clustering pattern of CP gene sequence in NN analysis was largely analogous to NJ phylogenetic tree, extensive networking within and among various CP gene groups was evident in NN analysis (Figure 4).

On the other hand, RdRp sequences corresponding to S5 and S9 phylogenetically clustered with well-characterized CTV genotype VT ‘*western*’ lineage sequences from different countries, while maintaining considerable distance from the unique VT sequence isolate (Kpg3; HM573451) from another part of NE India (Figure 5). These results largely substantiated with BLASTn results. Notably, NN reconstructed with RdRp sequences was substantially simpler, having scarcer evidences of inter and intragenotype networks (Figure 6), sharply contrasting the CP gene NN (Figure 4).

**Figure 3.**
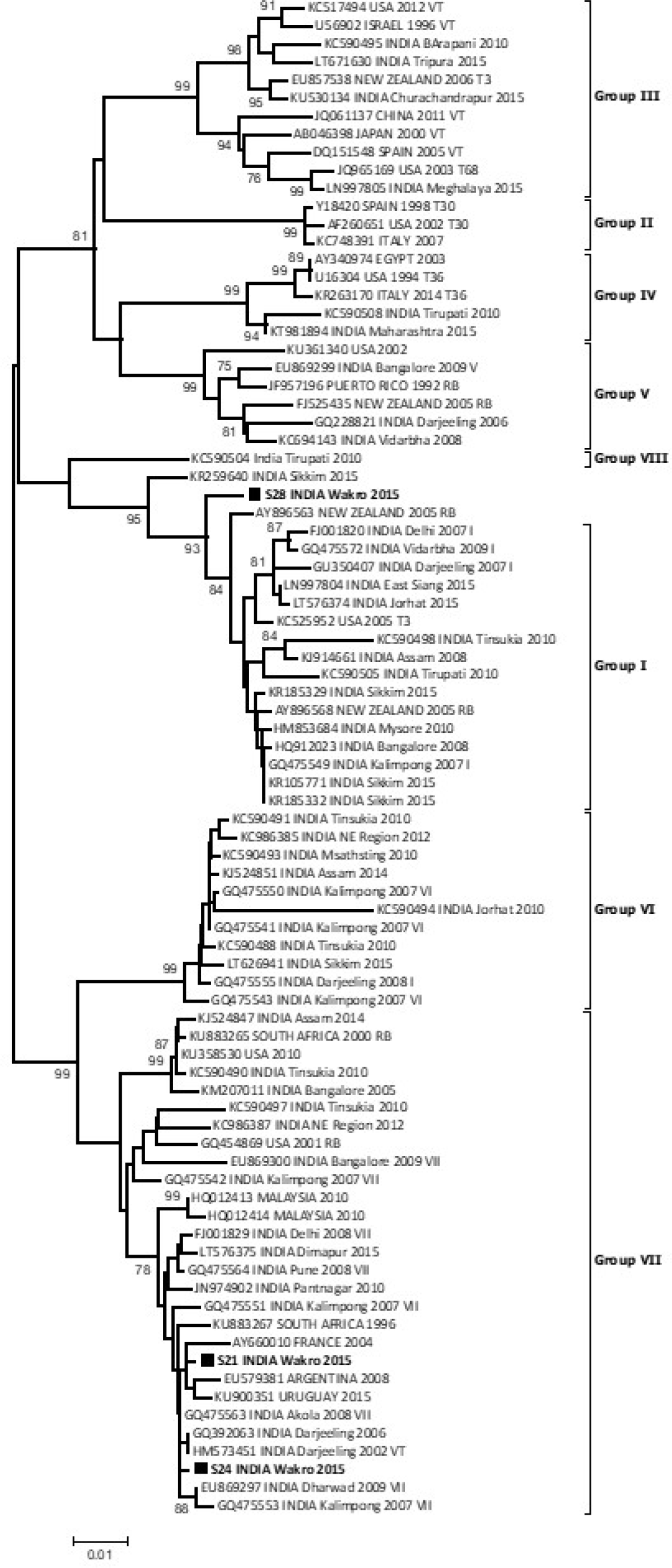
Phylogenetic tree reconstructed for CP (coat protein) gene sequences generated in the present study along with 81 other GenBank sequences, submitted by other researchers. Sequences from the present study (S21, S24 & S28) are marked by black solid squares. CP clades were classified as groups I to VIII following a previous literature (*Tarafdar et al*., *2013*). Evolutionary distances were calculated using the Maximum Composite Likelihood (MCL) method and phylogenetic relatedness was recreated using the Neighbor-Joining (NJ) method. Percentage of bootstrap support for each cluster (>75%) is shown at nodes.

**Figure 4.**
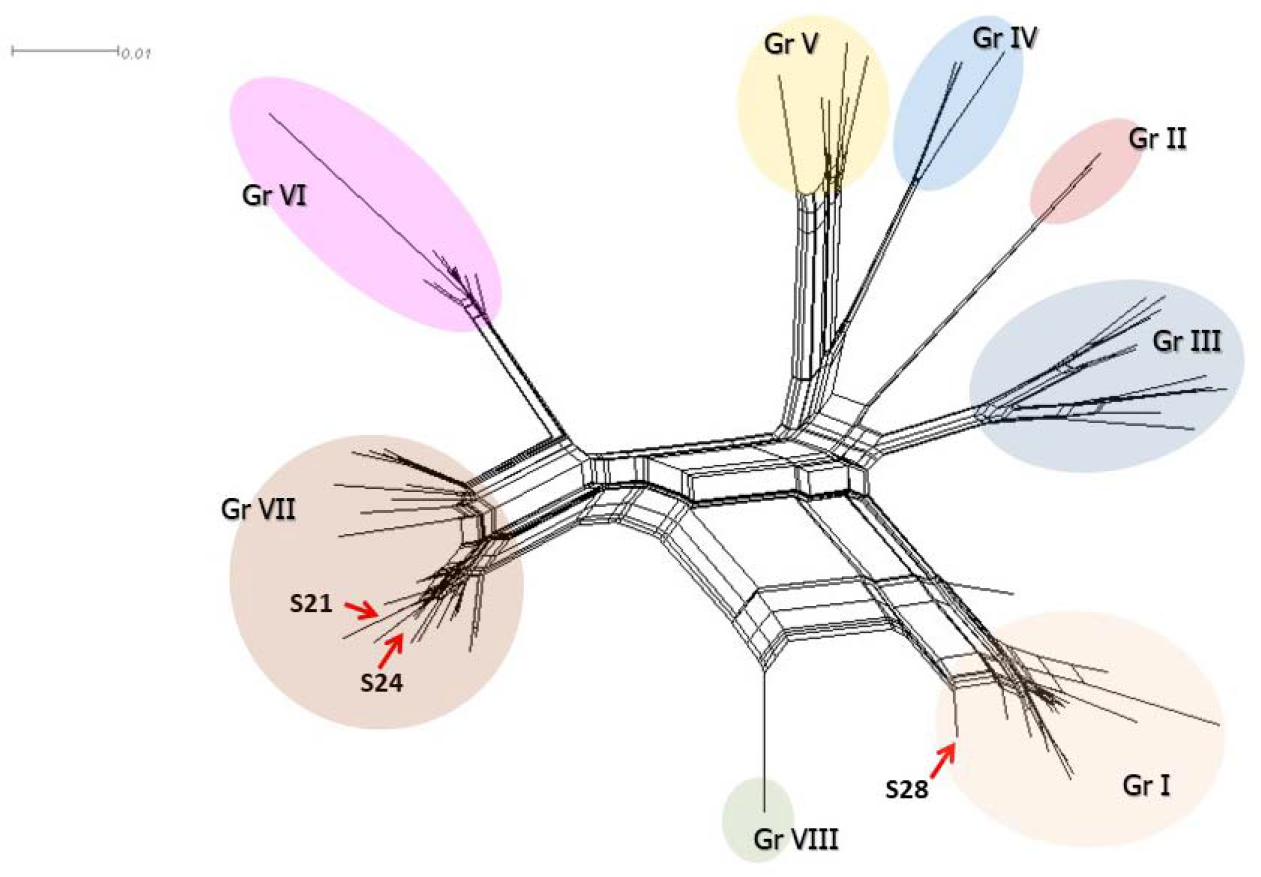
Neighbour Net of CP gene sequences reconstructed using the dataset previously used for reconstruction of NJ tree in Figure 3. Position of the three CP gene sequence generated in the present study (S21, S24 & S28) is indicated by red arrowheads.

**Figure 5.**
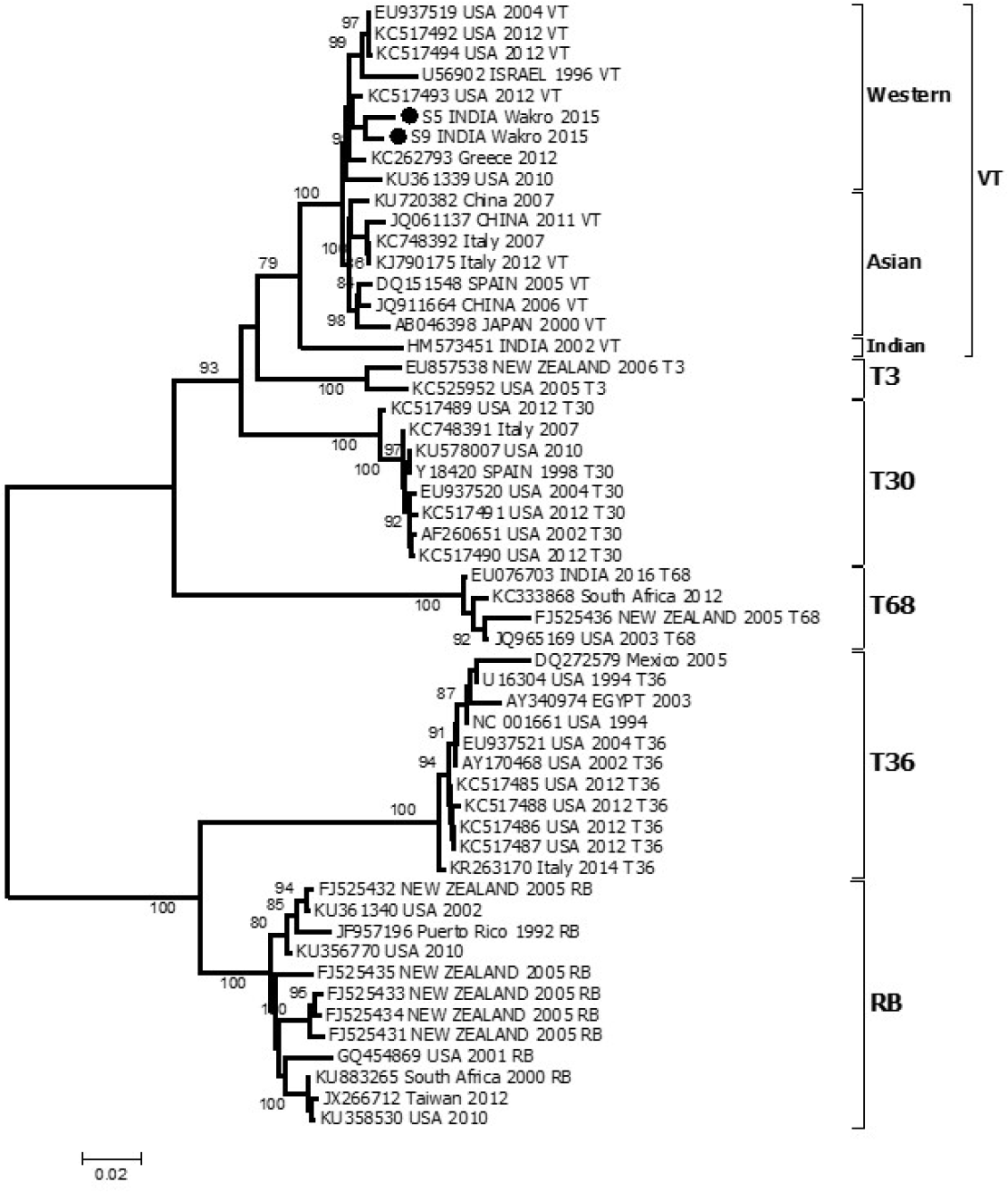
Phylogenetic tree reconstructed from 1860 bases of 5’ half of CTV genome (corresponding to partial ORF1a & ORF1b coding regions) from two sequence generated in this study (S5 & S9) and 52 others well characterized complete genomes retrieved from the GenBank. Sequences from the present study are marked by black solid rounds. Evolutionary distances were calculated using the Maximum Composite Likelihood (MCL) method and phylogenetic relatedness was recreated using the Neighbor-Joining (NJ) method. Percentage of bootstrap support for each cluster (>75%) is shown at the respective node. Genotypes and lineages are mentioned on the right-hand side.

**Figure 6.**
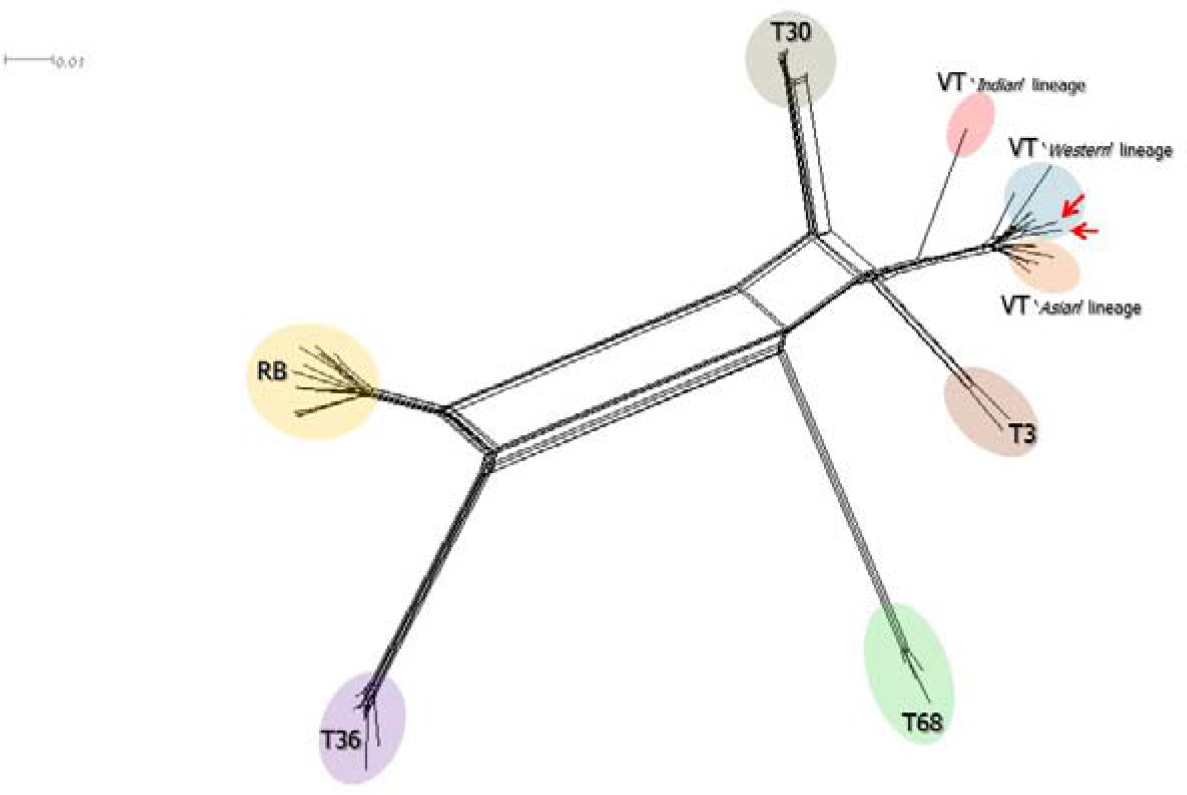
Neighbour Net reconstructed from 1860 bases of 5’ half of CTV genome (corresponding to partial ORF1a & ORF1b coding regions) using the dataset previously used for reconstruction of NJ tree in Figure 5. Position of the two-sequence generated in the present study (S5 & S9) is indicated by red arrowheads.

### Analyses of recombination

Among the two RdRp sequences generated in this study, sequence S5 had evidence of recombination (detected by Bootscan, MaxChi, SiScan and 3Seq with a probability of 1.346×10^−02^, 3.033×10^−06^, 2.166×10^−06^ and 1.510×10^−02^, respectively) between the present sequence S9 (minor parent) and a sequence resembling an isolate from China (major parent JQ061137, belonging to the ‘*Asian’* lineage of genotype VT) (Figure 7a). Interestingly, evidence of sequence S9 as the major parent was detected in all the four reference sequences belonging to genotype T68 (EU076703, India; FJ525436, New Zealand; JQ965169, USA and KC333868, South Africa), while as minor parent in both the reference sequences belonging to genotype T3 (EU857538, New Zealand and KC525952, USA) included in the present analysis.

On the other hand, analysing the CP gene sequence dataset, RDP program could not detect any evidence of recombination within sequences S21 and S24. However, isolate S28 was found to have a sequence fragment which had evidence of recombination (detected by SiScan and 3Seq with a probability of 1.243×10^−10^ and 1.336×10^−02^, respectively) between a sequence isolated from Reunion Island (major parent, AY660010, belonging to group VII) and a sequence (isolate *AG28*) from Tinsukia, NE India (minor parent, KC590498, belonging to group I) (Figure 7b). Nevertheless, considering the remote origin of sequence AY660010, and its intimate phylogenetic relatedness with sequence S21 (Figure 3), it is logical to interpret that an immediate ancestor (probably originating in NE India) of these two sequences might have actually been the major parent.

**Figure 7.**
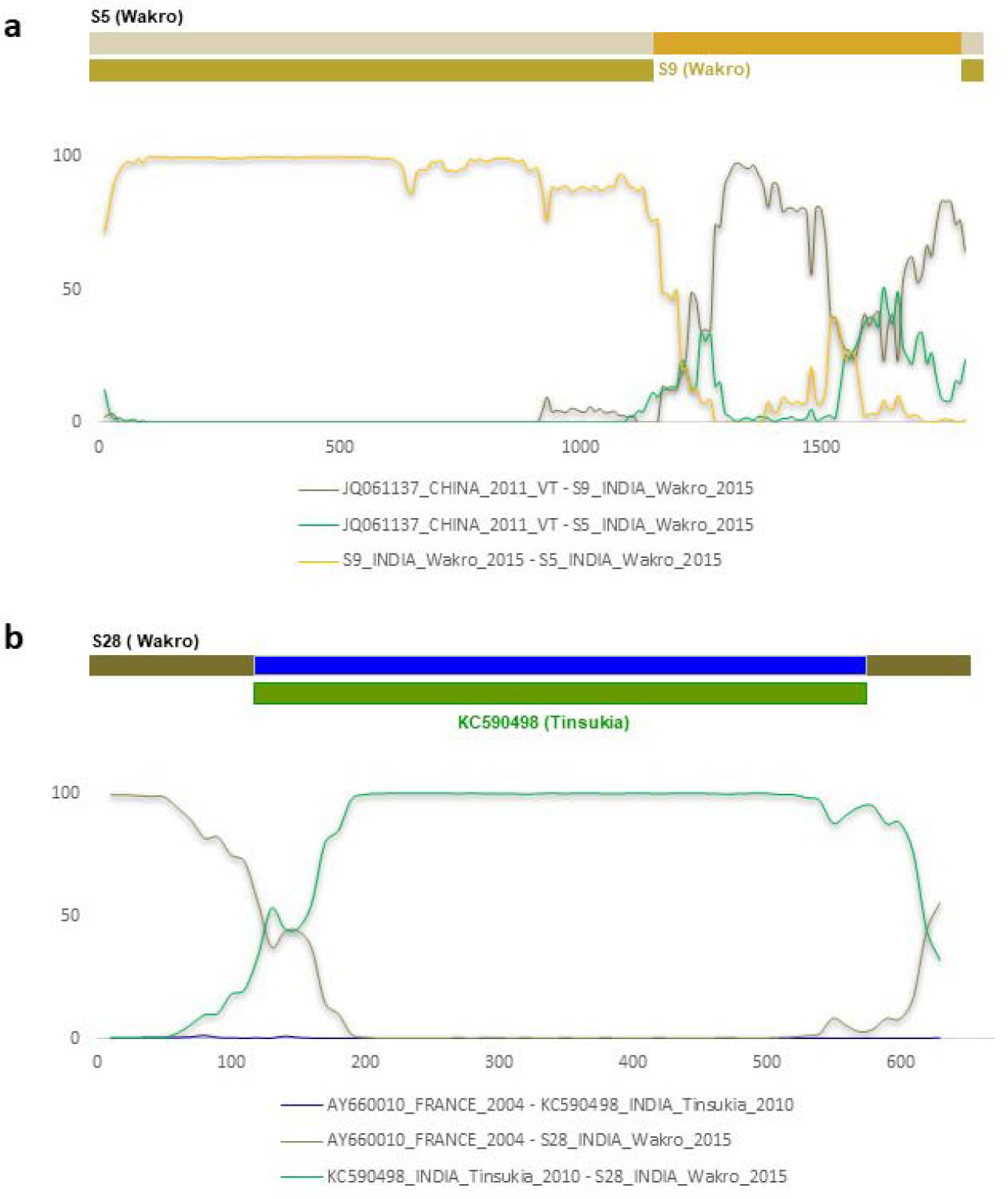
Recombination patterns observed in the sequences S5 (a) and S28 (b), generated in the present study. Each figure is followed by BootScan plot showing the breakpoints corresponding to the recombination event. Plots represent % of permuted trees on the *y* axis.

## DISCUSSION

The NE Indian biogeographic zone is an important part of the Indo-Burma biodiversity hotspot region, where the great Himalayan Mountains meets the Peninsular India. NE India comes under the *Hindustan (Indian) centre of origin of cultivated plant*, proposed by Nikolai Vavilov, and is considered the native place for numerous tropical and sub-tropical fruits including citrus^37,38^. It is also proposed that CTV has coevolved with its citrus hosts^2,38^. CTV is now ubiquitous and is the most devastating pathogen of citrus^2^. Similar to other citrus-growing regions of the world, CTV is highly prevalent in sub Himalayan regions of NE India, where it has severely affected and wiped-out several orchards^7,39^. Nevertheless, our understanding of CTV epidemiology in NE India is mostly restricted to accessible orchards in States of Assam, Meghalaya and Sikkim. To the best of our knowledge, there is no reliable literature on molecular epidemiology of CTV in the state of Arunachal Pradesh and this study is the first to report CTV molecular epidemiology in one of the oldest and remotest citrus-growing regions of India.

Previous studies have demonstrated an evolutionary asymmetry between the 5’and 3’ halves of CTV genome, where the 5’ (including the ORF1) is highly conserved and encodes genotype-specific phylogenetic signals while the 3’ (including the CP) shows relatively high genetic diversity^2,40,41^. Based on these observations, a ∼400 nucleotide region of ORF1a (corresponding to partial P-Pro domain) has been widely used for determining CTV genotypes^7,47^. In the present study, we amplified and analysed a comparatively longer genetic fragment (1860 bases corresponding to partial ORF1a & ORF1b coding regions) and found that phylogeny reconstructed with this genetic region is highly concordant with phylogenies derived from CTV complete genome sequences. Nevertheless, analyses of this genetic region in the present isolates demonstrated the presence of virulent genotype VT, substantiating with previous studies from the Indian subcontinent^7,14-20,36,42,^. Interestingly however, present isolates clustered with the ‘w*estern*’ lineage clade of VT sequences in the phylogeny. Since Wakro region geo-physically connects NE Indian part of Indian subcontinent with China and other parts of Southeast Asia, phylogenetic affiliation of present sequences to either ‘*Indian*’ or ‘*Asian*’ lineage was therefore highly anticipated. The *Western* and the *Asian* lineages signify evolution and global dispersal of two distinct variants of genotype VT in Western and Asian countries respectively^2^, the *Indian* lineage has been proposed to accommodate the decline inducing recombinant strain ‘*Kpg3*’ (GenBank accession HM573451), which do not cluster within either of the two lineages in phylogenies^2,7^.

Among extant CTV genotypes distributed globally, highest genetic diversity and widest geographical distribution of VT advocates its ancient origin. Analyses based on mixed effects model of evolution (MEME) of the *ORF1a*, further strengthens earliest origin of this genotype, providing additional cue that *western* lineage of VT is the most ancestral type known so far, from which all other lineages, including the *Asian* lineage have evolved through ‘*episodic diversification events*’^2^. Considering this geographical region as the proposed centre of origin of CTV, *Western* lineage CTV sequences found in present study appears to represent the original ‘*relict*’ CTV genotype. In such a scenario, natural spatio-temporal dispersal of this original virus to southeast Asia and peninsular India might have resulted in evolution of *Asian* and *Indian* lineages, respectively. In contrast, phylogenetic clustering of present sequences with genotype VT sequences from western countries amply support its relatively recent introduction to the western countries. Earlier, it has been proposed that CTV evolved and diverged into major clades in this region followed by their dispersal across Southeast Asia, Indian subcontinent and islands of Indian Ocean long ago, while their dissemination to Europe, Africa and Americas took place only recently (∼400-500 years ago)^3,6,44^.

Additionally, owing to stability of CTV genomes (evolutionary rates being the slowest among RNA viruses), similarity of genome sequences in distant geographical regions have been suggested to indicate origin and dispersal of CTV^3,6,44^. Interestingly, one of the ORF1 sequences (S5) in this study was found to be recombinant, involving a major parent similar to *Asian* lineage genotype VT sequence (JQ061137, China) and present isolate S9 sequence as minor parent. This finding of an *Asian* lineage sequence as major recombinant partner may either be suggestive of remnants of the *‘Asian*’ lineage evolved in this geographical region or might be a relatively recent introduction, which cannot be concluded from the present results. Notably, in recombination analysis, present sequence S9 was found to be involved in all the genotype T68 and genotype T3 reference sequence included in the present study.

On the other hand, previous studies on CP gene sequences have revealed circulation of diverse phylogroups in different citrus-growing regions, which do not necessarily correlate with the genotype determined by analysis of ORF1 sequences^17,38,45^. Such incongruence between different genetic regions of CTV genome has been attributed to frequent recombination events or gene flow among distinct genetic types occurring particularly in the 3’ half of the CTV genome^7,18,19,38,45^. Our findings of NN analyses of ORF1 and CP genetic regions showing sharply discrete pattern of reticulate evolution strongly support these previous observations.

Nevertheless, analysing CP gene variability, previous studies have reported eight globally circulating CP genetic variants (genogroups I to VIII), of which seven (I, III, IV, V, VI, VII, and VIII) are present throughout Indian subcontinent while five (I, III, V, VI and VII) are present in different parts of NE India^7,36,43^. In contrast, CP gene sequence diversity observed in present samples was limited to only two genogroups (I & VII), which were related to sequences from other parts of sub Himalayan NE India (Darjeeling, Kalimpong and Sikkim). We also detected a recombinant CP sequence involving a group VII sequence (resembling present sequence S21) accepting a large fragment from a group I sequence (similar to isolate AG28, GenBank KC590498) reported from Tinsukia, Assam^7,39^. However, we could not identify any genogroup VI CP gene sequence, which represents a clade of phylogenetically related sequences exclusively from different parts of NE India. Taken together, limited genetic diversity, evidence of recombination with sequences only from nearby geographical regions and remnants of the present sequences in other genotypes indicate the antiquity of present CTV sequences from Wakro and provides credible evidence to support prevailing hypotheses on origin of CTV.

Apart from CTV genotype, transmitting vector aphid species can also significantly influence the spatio-temporal dynamics and magnitude of infection. Although, BrCA (*T. citricida*) is the most frequent and efficient vectors associated with CTV transmission in India^13,15^, other aphid vectors (*T. aurantii, A. gossypii, Myzus persicae, Dactynotus jaceae, A. craccivora*) have also been recognized to transmit CTV^46^ During present survey, we could detect *T. aurantii* in high densities in all the declining orchards (situated at an altitude of 330-426 m). These findings are different from a previous study from Darjeeling, where *T. aurantii* was dominant in the citrus orchards at high altitude (>500 m), whereas *T. citricida* was predominant in the orchards of lower altitude^47^. *T. aurantii* has earlier been shown to be highly adaptable to wider host range and higher altitudes, particularly in the sub-tropical hills of NE India^48,49^. Remarkably, we could reproducibly amplify CTV specific 132 bp 3’-UTR genetic region in aphid pools, we could not amplify other genetic fragments (CP or the ORF1) from the aphid pools to implicate their role in transmission. Even though exactly similar observation of amplification of smaller viral RNA segments (130-162 nts) while non-amplification of larger segments^50^ have been reported from mosquito pools, reason for such phenomenon remains to be investigated.

In conclusion, present study reveals the molecular epidemiology of CTV from a place which comes under a geographical region, believed to be the origin of the citrus fruits. Due to its remoteness, this region has remained largely unreached and thus maintains pristine conditions in terms of biodiversity. Our results suggesting the circulation of an unanticipated genetic type of CTV with less interaction with CTV sequences only from neighbouring geographical areas indicate relict nature of present CTV sequences. However, we acknowledge that complete genome sequence would be required for better understanding the evolution and diversification of CTV.

## ACKNOWLEDGEMENT

This research was supported by institutional intramural grants from the Defence Research & Development Organization (DRDO), Ministry of Defence, Government of India. The authors gracefully thank Dr. BM Mishra, Ex-Deputy Commissioner (Lohit District), Government of Arunachal Pradesh, officials of the Horticulture department (AP) and the orange growers of Wakro for their support and co-operation during field survey.

## AUTHOR CONTRIBUTION

S.D. and V.V designed the study. S.D., R.G., V.M., M.K.M & S.R. performed field surveys, collected samples and performed identifications. S.D., B.D., R.B. performed laboratory experiments. S.D. analysed data, wrote the main manuscript, prepared the figures and tables. S.K.D. & V.V. overall supervised the research and critically reviewed the manuscript. All the authors except R.B. (deceased) reviewed the manuscript.

## REFERENCES

1. FAO. Citrus fruit fresh and processed statistical bulletin. http://www.fao.org/economic/est/est-commodities/citrus-fruit/en/ (2016).

2. Harper, S. J. Citrus tristeza virus: evolution of complex and varied genotypic groups. Front. Microbiol. 4, 93 (2013).

3. Moreno, P., Ambrós, S., Albiach-Martí, M.R., Guerri, J. & Peña, L. Citrus tristeza virus: a pathogen that changed the course of the citrus industry. Mol. Plant. Pathol. 9: 251–268 (2008).

4. Bar-Joseph M., Dawson, W. O. Citrus tristeza virus (eds. Mahy, B.W.J. & Van Regenmortel, M.H.V.) In Encyclopaedia Virology 3rd edition 1, 520–525 (Academic Press, 2008).

5. Lee, R.F. & Bar-Joseph, M. Tristeza. In Compendium of citrus diseases, 2nd edition 61–63 (APS Press, 2000).

6. Albiach-Martí, M.R., Mawassi, M., Gowda, S., Satyanarayana, T., Hilf, M.E., Shanker, S., et al. Sequences of citrus tristeza virus separated in time and space are essentially identical. J. Virol. 74, 6856–6865 (2000).

7. Biswas, K.K., Palchoudhury, S. & Ghosh, D.K. Closterovirus in India: Distribution, Genomics and Genetic Diversity of Citrus Tristeza Virus (eds. Mandal, B., et al.). In A Century of Plant Virology in India (Springer Nature, 2017).

8. Martelli, G. P., Agranovsky, A. A., Bar-Joseph, M., Boscia, D., Candresse, T., Coutts R. H. A., et al. Family Closteroviridae. In Ninth Report of the International Committee on Taxonomy of Viruses, Andrew, M.Q.K., Adams, M.J., Carstens, E.B., Leftkowitc, E.J., pp. 987–1001. (Elsevier Academic Press, 2012).

9. Kleynhans, J., Pietersen, G. Comparison of multiple viral population characterization methods on a candidate cross-protection Citrus tristeza virus (CTV) source. J. Virol. Methods. 237, 92–100 (2016).

10. ICAR-All India Coordinated Research Project on Fruits 2016-17. https://icar.org.in/dare-icar-annual-reports (2017).

11. Barbhuiya, A.R., Khan, M.L. & Dayanandan, S. Genetic structure and diversity of natural and domesticated populations of Citrus medica L. in the Eastern Himalayan region of Northeast India. Ecol. Evol. 6: 3898–3911 (2016).

12. Hynniewta, M., Malik, S.K. & Rao, S.R. Genetic diversity and phylogenetic analysis of Citrus (L) from north-east India as revealed by meiosis, and molecular analysis of internal transcribed spacer region of rDNA. Meta. Gene. 2, 237–251 (2014).

13. Ahlawat, Y.S. Viruses, greening bacterium and viroids associated with citrus (Citrus species) decline in India. Indian J. Agric. Sci. 67, 51–57 (1997).

14. Biswas, K.K. Molecular characterization of Citrus tristeza virus isolates from the Northeastern Himalayan region of India. Arch. Virol. 155, 959–63 (2010).

15. Biswas, K.K. Molecular diagnosis of Citrus tristeza virus in mandarin (Citrus reticulata) orchards of hills of West Bengal. Indian J. Virol. 19: 26–31 (2008).

16. Biswas, K.K., Tarafdar, A., Sharma, S. K., Singh, J. K., Dwivedi, S., Biswas, K., et al. Current status of Citrus tristeza virus incidence and its spatial distribution in citrus growing geographical zones of India. Indian J. Agri. Sci. 84: 8–13 (2014).

17. Biswas, K.K., Tarafdar, A., Diwedi, S. & Lee, R. F. Distribution, genetic diversity and recombination analysis of Citrus tristeza virus of India. Virus Genes 45, 139–48 (2012).

18. Biswas, K.K., Tarafdar, A., Sharma, S. K. Complete genome of mandarin decline Citrus tristeza virus of Northeastern Himalayan hill region of India: comparative analyses determine recombinant. Arch. Virol. 157: 579–83 (2012).

19. Sharma, S.K., Tarafdar, A., Khatun, D., Kumari, S. & Biswas, K.K. Intrafarm diversity and evidence of genetic recombination of Citrus tristeza virus isolates in Delhi region of India. J. Plant Biochem. Biotechnol. 21: 38–43 (2012).

20. Ghosh, D. K., Aghave, B., Roy, A. & Ahlawat, Y.S. Molecular cloning, sequencing and phylogenetic analysis of coat protein gene of a biologically distinct CTV isolate occurring in central India. J. Plant Biochem. Biotechnol. 18, 105–108 (2009).

21. Geographical Indications Journal, volume 61, Geographical Indications Registry, Government of India. November 21, 2014. http://www.ipindia.nic.in/journal-gi.htm

22. Business Standard. Orange growth hit in Arunachal Pradesh. January 11, 2015. https://www.business-standard.com/article/pti-stories/orange-growth-hit-in-arunachal-pradesh-115011100455_1.html

23. Arunachal Govt portal; Citrus decline is alarming. Published on State Portal of Arunachal Pradesh http://arunachalpradesh.nic.in/csp_ap_portal/pdf/Documents/citrus-decline-alarming-arunachal.pdf)

24. The Assam Tribune; Citrus decline hits orange bowl of Arunachal. Guwahati, Monday, January 19, 2015. http://www.assamtribune.com/scripts/mdetails.asp?id=jan1915/oth052

25. ICAR-NEH Annual Report 2014-15. ICAR Research Complex for NEH Region Umroi Road, Umiam, Meghalaya. www.icarneh.ernet.in and http://kiran.nic.in

26. Olmos, A., Cambra, M., Esteban, O., Gorris, M. T. & Terrada, E. New device and method for capture, reverse transcription and nested PCR in a single closed-tube. Nucleic Acids Res. 27, 1564–1565 (1999).

27. Huang, Z., Rundell, P.A., Guan, X. & Powell, C.A. Detection and Isolate Differentiation of Citrus tristeza virus in infected trees based on Reverse Transcription-Polymerase Chain Reaction. Plant Dis. 88, 625–629 (2004).

28. Vives, M.C., Rubio, L., Sambade, A., Mirkov, T.E., Moreno, P. & Guerri, J. Evidence of multiple recombination events between two RNA sequence variants within a Citrus tristeza virus isolate. Virology. 331, 232–237 (2005).

29. Hall, T. A. BioEdit: a user-friendly biological sequence alignment editor and analysis program for windows 95/98/NT. Nucleic Acids Symp. Ser. 41, 95–98 (1999).

30. Altschul, S.F., Gish, W., Miller, W., Myers, E.W. & Lipman, D.J. Basic local alignment search tool. J. Mol. Biol. 215, 403–410 (1990).

31. Edgar, R.C. MUSCLE: multiple sequence alignment with high accuracy and high throughput. Nucleic Acids Res. 32, 1792–1797 (2004).

32. Tamura, K., Stecher, G., Peterson, D., Filipski, A., Kumar, S. MEGA 6: molecular evolutionary genetics analysis version 6.0. Mol. Biol. Evol. 30, 2725–2729 (2013).

33. Huson, D.H. & Bryant, D. Application of Phylogenetic Networks in Evolutionary Studies. Mol. Biol. Evol. 23, 254–267 (2006).

34. Woolley, S. M., Posada, D. & Crandall, K.A. A comparison of phylogenetic network methods using computer simulation. PLoS One 3: e1913 (2008).

35. Martin, D.P., Murrell, B., Golden, M., Khoosal, A. & Muhire, B. RDP4: Detection and analysis of recombination patterns in virus genomes. Virus Evol. 1, vev003 (2015).

36. Tarafdar, A., Godara, S., Dwivedi, S., Jayakumar, B.K. & Biswas, K.K. Characterization of Citrus tristeza virus and determination of genetic variability in North-east and South India et al. Indian Phytopath. 2013, 66, 302–307 (2013).

37. Hummer, K.E. & Hancock, J.F. Vavilovian Centers of Plant Diversity: Implications and Impacts. Hortscience. 50, 780–783 (2015).

38. Martin, S., Sambade, A., Rubio, L., Vives, M. C., Moya, P., Guerri, P., Elena, S. F. & Moreno, P. Contribution of recombination and selection to molecular evolution of Citrus tristeza virus. J. Gen. Virol. 90, 1527–1538 (2009).

39. Biswas, K.K., Pal Choudhuri, S. & Godara, S. Decline of Mandarin Orange Caused by Citrus tristeza virus in Northeast India: Conventional and Biotechnological Management Approaches. J. Agri. Eng. Food Tech. 3, 236–241 (2016).

40. Hilf, M.E., Mavrodieva, V.A., Garnsey, S.M. Genetic marker analysis of a global collection of isolates of Citrus tristeza virus: characterization and distribution of CTV genotypes and association with symptoms. Phytopathology. 95, 909–917 (2005).

41. Roy, A., Brlansky, R.H. Genome analysis of an orange stem pitting citrus tristeza virus isolate reveals a novel recombinant genotype. Virus Res. 151, 118–130 (2010).

42. Singh, J.K., Tarafdar, A., Sharma, S.K. & Biswas, K.K. Evidence of Recombinant Citrus tristeza virus Isolate Occurring in Acid Lime cv. Pant Lemon Orchard in Uttarakhand Terai Region of Northern Himalaya in India. Indian J. Virol. 24, 35–41 (2013).

43. Palchoudhury, S., Ghimiray, S., Biswas, M.K. & Biswas, K.K. Citrus tristeza virus Variants and their Distribution in Mandarin Orchards in Northeastern Himalayan Hill Region of India. Int. J. Curr. Microbiol. App. Sci. 6: 1680–1690 (2017).

44. Silva, G., Marques, N. & Nolasco, G. The evolutionary rate of citrus tristeza virus ranks among the rates of the slowest RNA viruses. J. Gen. Virol. 93, 419–429 (2012).

45. Rubio, L., Ayllón, M. A., Kong, P., Fernández, A., Polek, M., Guerri, J., Moreno, P. & Falk, B.W. Genetic variation of Citrus tristeza virus isolates from California and Spain: evidence for mixed infections and recombination. J. Virol. 75, 8054–8062 (2001).

46. Verma, P.M., Rao, D.G. & Capoor, S.P. Transmission of tristeza virus by Aphis craccivora Koch and Dactynotus jaceae (L.) Indian J. Entomol. 27, 67–71 (1965).

47. Ghosh, A., Das, A., Lepcha, R., Majumdar, K. & Baranwal, B.K. Identification and distribution of aphid vectors spreading Citrus tristeza virus in Darjeeling hills and Dooars of India. J. Asia Pacific Entomol. 18, 601–605 (2015).

48. Agarwala, B.K. & Bhattacharya, S. Anholocycly in tropical aphids: population trends and influence of temperature on development, reproduction, and survival of three aphid species (Homoptera: Aphidoidea). Phytophaga 6, 17–27 (1994).

49. Roychoudhuri, D.N. Aphids of North-east India and Bhutan. pp. 76–78 (Zoological Society of India, 1980).

50. Ravanini, P., Huhtamo, E., Ilaria, V., Crobu, M.G., Nicosia, A.M., Servino, L., et al. Japanese encephalitis virus RNA detected in Culex pipiens mosquitoes in Italy. Euro. Surveillance 17, 20221 (2012).

